# Spatial risk of disease transmission between wild bovids and livestock in Thailand

**DOI:** 10.1101/2024.05.04.592526

**Authors:** Wantida Horpiencharoen, Jonathan C. Marshall, Renata L. Muylaert, Reju Sam John, David T. S. Hayman

## Abstract

The livestock-wildlife interface is one of the most essential issues threatening wildlife conservation and public health. Identifying interface areas can help to prioritise disease surveillance and implement mitigation measures and control programs for targeting threatened wildlife. We predicted interface areas which were assumed to be areas at risk of infectious disease transmission based on the spatial overlap between three Thai wild bovids (including gaur, banteng and wild water buffalo) habitat suitability and domestic cattle. We assumed that domestic cattle are the reservoir of bovine infectious disease, and that high cattle density is a proxy for a higher risk of disease transmission. Our study indicated that the highest risk areas for the native species are at the forest edges where overlap exists between high habitat suitability and high cattle density. Wild water buffalo showed the largest proportion of high-risk areas (8%), while gaur and banteng showed similar risk areas (4%) in Thailand. The largest proportion of risk areas overlapping with protected areas was Namtok Sam Lan PAs at 89% for gaur, 84% for banteng and 65% for wild water buffalo. Kuiburi NP has the largest risk area around 274 km^2^ (around 28% of the total protected area) for gaur and banteng, whereas wild water buffalo has the largest risk area overlapping with Huai Thabthan-Had Samran around 126 km^2^ (10% of the PA). Kaengkrachan Forest Complex showed the second largest risk area from 249 km^2^ for gaur and 273 km^2^ for banteng (8-9% of the PA). Our results address how habitat suitability might be helpful for infectious disease prevention and control strategies focused on native fauna and One Health. Furthermore, this work may also support the wild bovid habitat conservation initiatives and land use planning by informing decision-making about balancing wildlife habitats and livestock farming.

## Introduction

Wild Bovidae (Mammalia: Artiodactyla) are distributed worldwide and play crucial ecosystem roles, because they determine the forest and ecosystem structure, transport micronutrients, and disperse plant seeds (1, 2) and are also important prey species of predators (3). In Asia, wild bovid populations are threatened by multiple factors, including habitat loss and hunting, especially in South to Southeast Asia (4). Natural habitat loss often comes with increased free-grazing livestock interaction, which can lead to problems as varied as resource competition, reducing wildlife population abundance (5), interbreeding between domestic and wild water buffalo (6), and infectious disease transmission (7).

Among the twenty-seven recognized wild bovid species as of 2020 (IUCN), five species remain in Thailand, including gaur (*Bos gaurus*), banteng (*Bos javanicus*), wild water buffalo (*Bubalus arnee*), mainland serow (*Capricornis sumatraensis*) and Chinese goral (*Naemorhedus griseus*). Their habitat and populations have been threatened by human activities such as deforestation and hunting (8-10). Wild bovids, especially the large herbivores (e.g. gaur, banteng and wild water buffalo), gradually adapt their distribution and behaviour to land use change. For example, gaur has been found close to agricultural areas and forest edges where they forage on crop plantations (e.g. grass, cassava) (11). Banteng is also well-adapted to secondary forests near villages and logging sites (12). These wild bovids are, therefore, able to share natural resources with free-grazing domestic bovids, which can potentially cause disease transmission via direct and indirect contact with the sources of infections (e.g. infectious cattle, host, environmental reservoirs (13).

The livestock-wildlife interface is one important issue threatening wildlife conservation and global public health because 72% of reported emerging diseases originate from wildlife to humans and/or livestock (14). Bovine infectious diseases, such as bovine tuberculosis, brucellosis and foot and mouth disease, can be transmitted and circulate in domestic and wild bovid populations (15). These diseases and their impact on wildlife and livestock population health have been studied in Europe (16), North America (17) and Africa (18), but less so in Asia, including Thailand.

Several factors can drive disease transmission between livestock and wildlife populations, such as the expanding livestock production (19), the shrinking of wildlife habitat (20) and changes in wildlife distribution, demography and behaviour (21). Among these factors, high host density is potentially a determinant risk factor that can lead to successful disease transmission as it may translate to a higher probability of between- and among-host species interactions, contact and pathogen exposure (22, 23). The movement and spatial overlap of wildlife and livestock can lead to increased infectious disease transmission risk. Areas where there is potential for interaction between a new susceptible host and a reservoir can increase the chance of disease transmission through increasing contact rates and time (24). For wild and domesticated species, these areas are usually the transition areas between two or more land use types, such as the edges of forest and agricultural areas, which are likely to have more species activities leading to a greater chance of interaction and so disease transmission among wildlife and livestock (25). Previous studies indicated the presence of some infectious diseases, such as babesiosis and leptospirosis (26, 27) in domestic cattle at the edge of forest, making these domestic animals a potential reservoir of disease transmission to the wild bovids.

Bovine infectious diseases can circulate between livestock and wild ungulates with varying levels of virulence (15). Certain pathogens can circulate within either livestock or wildlife populations without causing visible clinical signs but have a significant impact on other species. For example, foot and mouth disease (FMD) might not affect African buffalo, but can lead to mortality in gaur (28). Similarly, haemorrhagic septicaemia (*Pasteurella multocida*) might be identified within the farm environment as non-pathogenic with limited mortality except under certain circumstances but can cause mass mortality in saiga (29). Chronic diseases like bovine tuberculosis and brucellosis with long incubation and relatively low fatality rates could potentially have long-term consequences by reducing populations due to disease, while FMD, which has a higher transmission rate with short incubation periods or even highly fatal infections such as anthrax, may have lower impacts on populations (30)

Moreover, in the past twenty years, there have been numerous transboundary emerging disease outbreaks among domestic animals in Thailand. For example, there have been outbreaks of lumpy skin disease among cattle (31), African horse sickness among horses (32), and African swine fever among pigs (33). Hence, it is crucial to investigate where there are high risk areas to prevent disease transmission to wild populations, considering their susceptibility to similar pathogens shared by livestock. Targeting the potential risks of disease transmission in wildlife and livestock interface areas can support the implementation of surveillance and control measures that may help prevent cross-species transmission (34).

In this study, we aim to 1) identify the potential risk of disease transmission of wild bovids and livestock in Thailand and 2) provide suggestions for disease surveillance and conflict mitigation measures in the wildlife-livestock interface areas of Thailand. The outputs could be used to prioritise local surveillance and mitigation measures for optimising resource allocation.

## Methods

### Study area

Thailand is located on the Indochina Peninsula, part of mainland Southeast Asia. Thailand borders four countries, Myanmar, Laos, Cambodia and Malaysia, with the Gulf of Thailand on the southeast connected to the Pacific Ocean and the southwest connected to the Andaman Sea. The total country area is around 514,000 km^2^, with agricultural land covering 41% and forested areas comprising around 32% of the country area. Most agricultural areas consist of rice fields (51%) and crop plantations (34%), while livestock farming covers only around 0.6% of the total agricultural area or around 0.3% of the total country (land use data source: https://agri-map-online.moac.go.th/). There is high cattle production in the central west, which includes free-range cattle and buffalo in some rural areas. These free-ranging livestock have encroached into wildlife habitats and share the same resources. Moreover, there is shared land use, for example, domestic buffalo may use rice fields.

### Populations

This study focuses on wildlife and livestock populations, and we calculate the largest potential for wildlife-livestock interface areas. We selected the remaining wild bovid species in Thailand because they are widely distributed and likely to share the same resources and pathogens as livestock, especially the large bovids (gaur, banteng and wild water buffalo) distribution, which tends to overlap with free-ranging cattle and agricultural areas. For the livestock population, we used the cattle population estimates as cattle production is all over the country, with varied production scales and systems from intensive farming to free grazing. We assume that domestic cattle can be a pathogen reservoir and transmit diseases to the wild bovid population, and our focus is on livestock transmitting infection to wild species, though the alternative is possible. Therefore, for our analysis, a high cattle density is assumed to have a higher risk transmission risk and a lower cattle density have a lower risk, as reported in previous studies (25, 35).

### Identifying the potential risk

Briefly, we identified the risk area using two types of datasets: 1) wild bovid distribution and 2) cattle density. Then, we overlaid these together and calculated the overlapping areas in 1-km^2^ cells as a sampling unit.

#### Wild bovid potential distribution

We assumed wild bovid distributions correlate with their suitable habitat we previously predicted by ecological niche models (36). Ecological niche models used 28 as predictor environmental variables using 8 algorithms. We conducted the ensemble models using the weighted mean method and used True Skill Statistics as a threshold to convert the ensemble models to binary values (1 = suitable areas and 0 = unsuitable). We selected three wild bovid species for further analysis: gaur (*B. gaurus*), banteng (*B. javanicus*), and wild water buffalo (*B. arnee*), and excluded Mainland serow (*C. sumatraensis*) or Chinese goral (*N. griseus*) from the analyses because the ecological niche models did not perform well. Full methods and model results can be found in Horpiencharoen et al. (2023) (36).

#### Cattle density

We downloaded cattle density data from Global Livestock of the World 2015, GLW 4 ([link](https://dataverse.harvard.edu/dataverse/glw_4;jsessionid=4b0179def6cd7ca0ec51b55e6444)). This data gives values of cattle density at an original spatial resolution of 10 km^2^. We cropped the raster layer to Thailand limits and disaggregated the raster to 1 km^2^ per cell to make it compatible with the habitat suitability raster using the raster package (37). Then, we rescaled the density values to 0 - 1 using this equation:

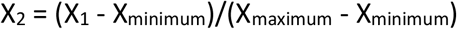

Where X is the value in the cattle density cell. Then, we used the mean of cattle density (0.14 cattle/km^2^) in Thailand calculated from the GLW 4 raster as a cut-off value for converting the cattle density raster into the binary values of high and low. The raster cells containing values greater than the mean were converted to 1 (high density), and the values lower than the mean were converted to 0 (low density).

We assumed that higher cattle density correlates with a greater risk of infectious diseases. Therefore, we counted the number of outbreaks in low and high cattle density areas to test this assumption. We divided the total number of outbreak events by the total area of cattle density for each category to check whether the higher number of outbreaks in high-density areas was not simply due to larger areas, as the following calculation:

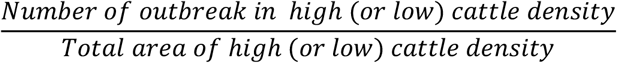

The results found a higher incidence of outbreak events in high cattle density areas compared to low cattle density areas. Thus, we used cattle density as the major risk factor to identify the potential disease transmission areas (more details in the results Table 2**Error! Reference source not found**.).

#### The potential risk areas

In this study, the potential high risk areas refer to the interface areas between wildlife and livestock that potentially share the resources (e.g. water bodies, grassland, mineral lick) and have a higher risk of disease transmission due to the increased opportunity of direct and indirect contact with disease reservoirs and environment, compared to other areas (25, 35, 38).

To define the risk areas, we overlapped the cattle density raster with the species’ binary maps and calculated the percentages of the potential risk areas in Thailand. Then, we intersected the risk areas with the national protected areas (PA) (39) to calculate the risk areas inside and outside PAs and also classified the risk areas by land use types to prioritise where to implement the disease surveillance. Lastly, we counted the occurrence of disease outbreaks reported by the Department of Livestock Development, Thailand, within the interface areas to explore the distribution of highly frequent diseases in the high risk areas (see below). All spatial analyses were programmed in R 4.3.1 (40). The code is available at a public repository (https://github.com/Wantidah/BovidRiskMaps). Data is available upon reasonable request.

#### Disease occurrence data

We used the national database of livestock disease outbreak reports from 2013 to 2021 generated by the Department of Livestock Development, Thailand. The data collection starts when an outbreak in livestock is reported by local authorities or farm owners.

Then epidemiological data are recorded, including the date of the index case, animal type, clinical signs, and number of animals infected, followed by collecting the samples for laboratory diagnosis. If a diagnosis is made and the causative pathogen and disease known, the authorities will record this in the database. If the authorities cannot find the causing pathogen, they will add the tentative diagnosis from the clinical signs. Each outbreak will be reported as confirmed if the causative pathogen is identified by laboratory diagnostics. However, if there is no laboratory result, the authority will fill in the tentative diagnosis according to animal clinical signs. The GPS location of the outbreak refers to the centroid of the sub-district (average area of districts of Thailand = 87 km^2^, range: 0.88 - 2,387 km^2^) where the outbreak occurred.

Here we selected five globally or regionally common bovine infectious diseases considered important for livestock health: 1) foot and mouth disease (FMD), 2) haemorrhagic septicaemia (HS - *Pasteurella multocida*), 3) bovine Tuberculosis (*Mycobacterium bovis* - bTB), 4) lumpy skin disease (LSD) and 5) brucellosis (*Brucella abortus*) from the national database. We selected outbreaks from these five diseases in cattle, then cleaned the outbreak events by excluding incorrect coordinates falling outside Thailand using R. Lastly, we counted the number of outbreaks within overlapping areas for each species and cattle population densities using the ‘extract’ function in the raster R package (37).

## Results

The high-risk areas with high wild bovid habitat suitability and high cattle density are mostly found in the central-western through the southern part of Thailand for the three species (Figure 1). The districts that showed the highest percentages of the risk areas are Nakhon Si Thammarat (south), Ratchaburi and Prachuap Khiri Khan) for all three species. Wild water buffalo showed the largest of the total interface areas, covering ~44,000 km^2^ (8% of Thailand), due to their potential habitat suitability predicted across the country. However, in the actual species distribution, only one population remains in the Huai Kha Khaeng Wildlife Sanctuary. Banteng and Guar showed similar potential habitat suitability species, which also resulted in the closest number of interface areas, ~22,000 km^2^ (4% of Thailand) (Table 1).

**Table 1.** The percentage of interface areas overlapped with protected areas by three wild bovid species.

| Species | Interface area with livestock (km<sup>2</sup>) |  |  | Percentage (%) of overlap area in Thailand |
| --- | --- | --- | --- | --- |
|  | inside PA | outside PA | total |  |
| Gaur | 2,018 | 18,367 | 20,385 | 3.9 |
| Banteng | 2,089 | 20,477 | 22,566 | 4.4 |
| Wild water buffalo | 747 | 43,642 | 44,389 | 8.7 |

**Table 2.** The occurrence of outbreak events classified by infectious disease and cattle density from 2013 to 2021.

| Infectious disease | Cattle density |  | Total |
| --- | --- | --- | --- |
|  | Low (<=mean) | High (>mean) |  |
| bTB | 2 | 0 | 2 |
| HS | 3 | 2 | 5 |
| LSD | 445 | 6,473 | 6,918 |
| FMD | 175 | 395 | 563 |
| Brucellosis | 15 | 24 | 39 |
| Total | 640 | 6,894 | 7,534 |
| Area of Thailand (km<sup>2</sup>) | 324,335 | 190,076 | 514,411 |
| Total outbreak event per area (km<sup>2</sup>) | 0.0020 | 0.0363 | 0.0146 |

**Figure 1.**
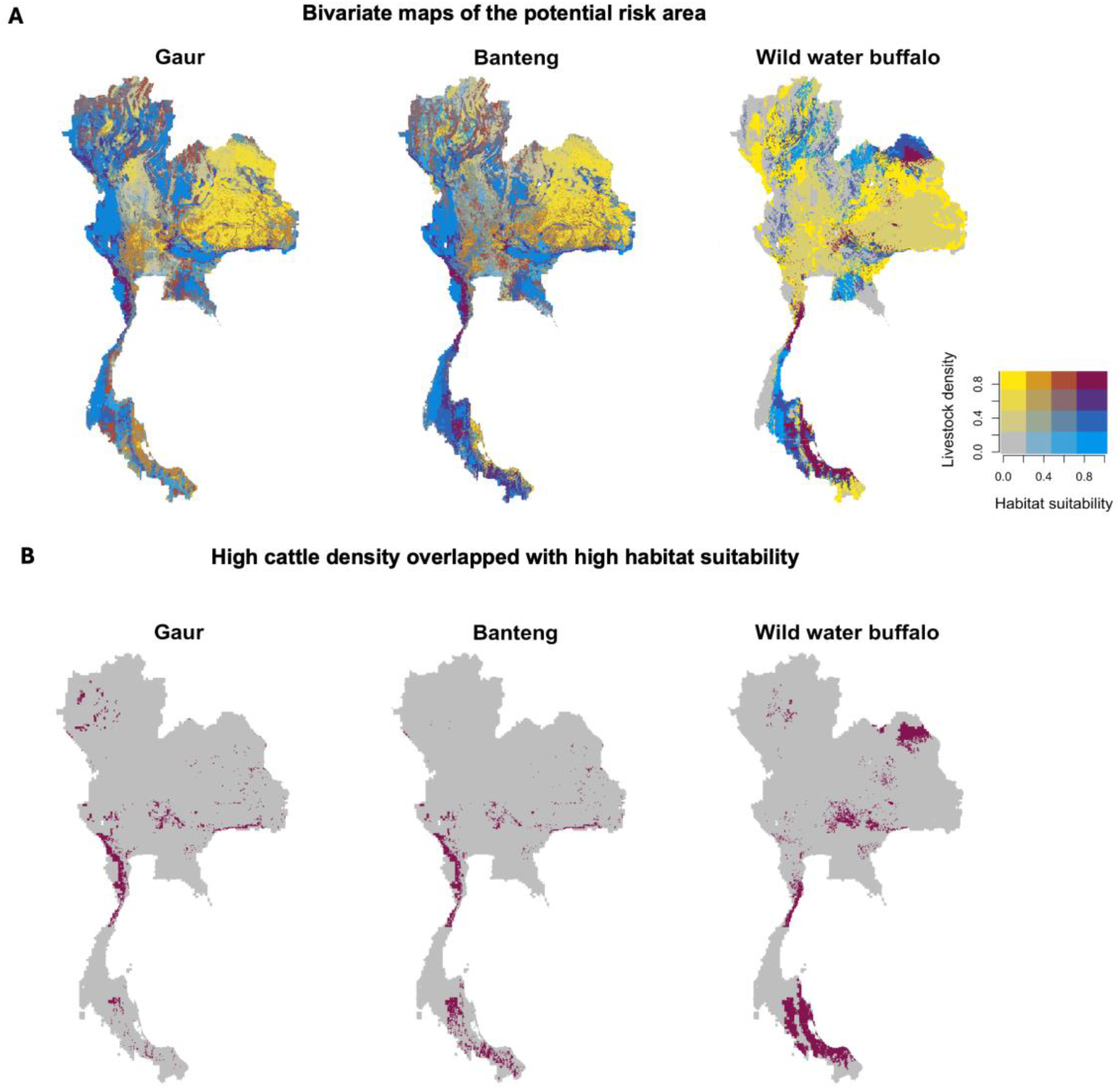
Bivariate maps of the potential risk area (A) between the habitat suitability (blue) and cattle density (yellow) reveal the interface areas between three wild bovid species and domestic cattle populations in Thailand. High-risk areas are represented in dark red, while low-risk areas are represented in grey. The potential high risk areas for disease transmission between three wild bovid species and cattle populations in Thailand (B). High-risk areas are represented in dark red, extracted from the interface areas in (A).

The highest percentage of risk areas inside the PA were identified in Namtok Sam Lan (also known as Phra Budda Chai) National Park (NP) (45 km^2^) in Saraburi Province, covering approximately 89% for gaur, 83% for banteng and 65% for buffalo. The second highest percentage for gaur and banteng is Namtok Huai Yang (160 km^2^) NP in Prachuap Khiri Khan, covering 60% (~100 km^2^) of the total PA, and for wild water buffalo is Huai Thabthan-Had Samran (498 km^2^) representing 25% (125 km^2^) of the total PA. However, for gaur and banteng, the largest risk area is located in the same PA—Kuiburi NP (970 km^2^), representing 273 km^2^ (28% of the PA). This is followed by the Kaengkrachan forest complex, representing 249 km^2^ for gaur (8% of the PA) and 261 km^2^ for banteng (9% of the PA). These two protected areas are in close proximity, with high-risk areas situated along the western forest edge, connected to agricultural areas with high cattle density, while the western side is connected to the Myanmar forest (Figure 1). Moreover, the large intact forests like the Western, Eastern and Dong Payayen - Khoa Yai forest complex illustrated high habitat suitability with low cattle density within the PA, but showed high risk at the border of the forests, while the fragmented forests in the north illustrated the potential high risk of disease transmission with high cattle density and low habitat suitability.

According to the national disease surveillance, the total number of outbreak events is 7,522 events for the five selected bovine infectious diseases from 2013 to 2021. LSD (6,913) has the most recorded outbreak events among the others, followed by FMD (563) and brucellosis (39), while HS and bTB have only 5 and 2 events, respectively (Table 2 and Figure 2). This is because the first LSD outbreak occurred in cattle herds in Thailand in 2021, leading to a large number of events reported across the country in a short period, while the other infections, which have lower records, are endemic in this area.

**Figure 2.**
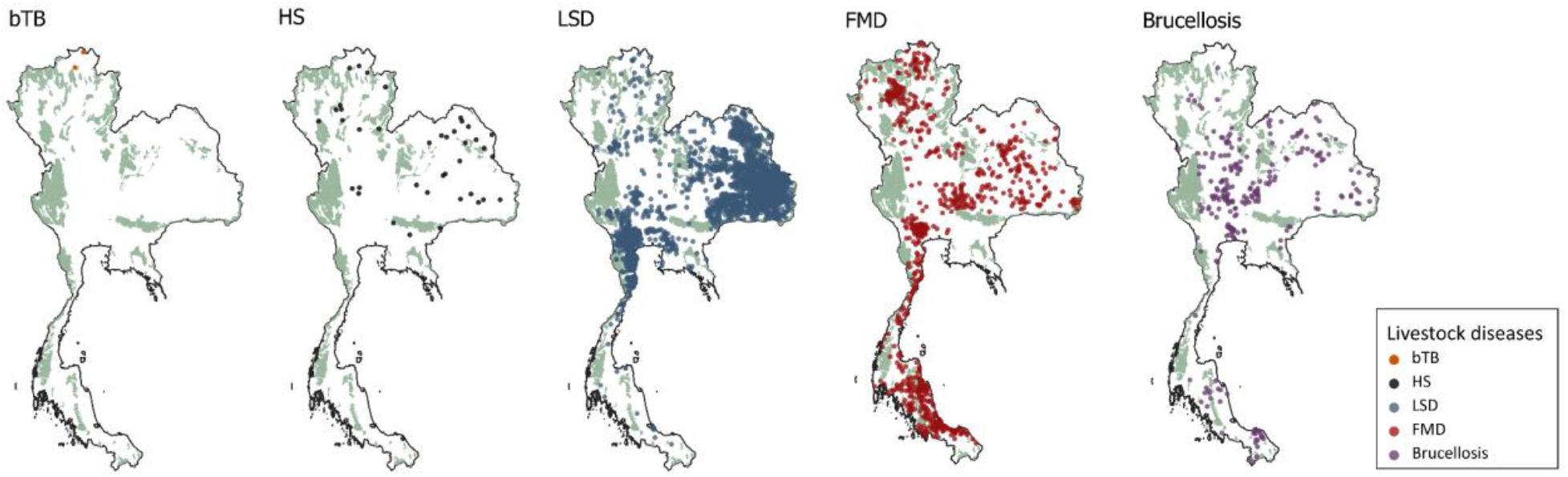
Occurrence of recorded bovine infectious disease outbreaks in Thailand from 2013 to 2021 and protected area distribution (PA; green). The maps show that the outbreaks occurred across Thailand, particularly in proximity to protected and forest areas that overlap with suitable habitats for wild bovids.

The cattle density demonstrated correlations with the number of infectious disease outbreaks, and this correlation is proportional to the area size. We observed that in high cattle density areas (190,076 km^2^), there were higher outbreak events, totalling around 6,894 events (0.036 events per km^2^), 18 times more than low cattle density areas (324,335 km^2^), which had 640 events (0.002 events per km^2^). The results of outbreak events by cattle density areas are presented in Table 2

We found that the density of outbreak events in cattle within the potential high risk area of gaur (0.01) and especially for wild water buffalo (0.015) were similar to the average density calculated for the country (0.0146) (Table 3). Wild water buffalo showed the highest events (647) within the risk areas as they have the largest potential habitat areas across the country while gaur (199) and banteng (162) show close results to each other. Similar to Table 2, the greatest numbers of disease events within high risk areas was for LSD and FMD in all species and the other diseases presented only small numbers.

**Table 3.**
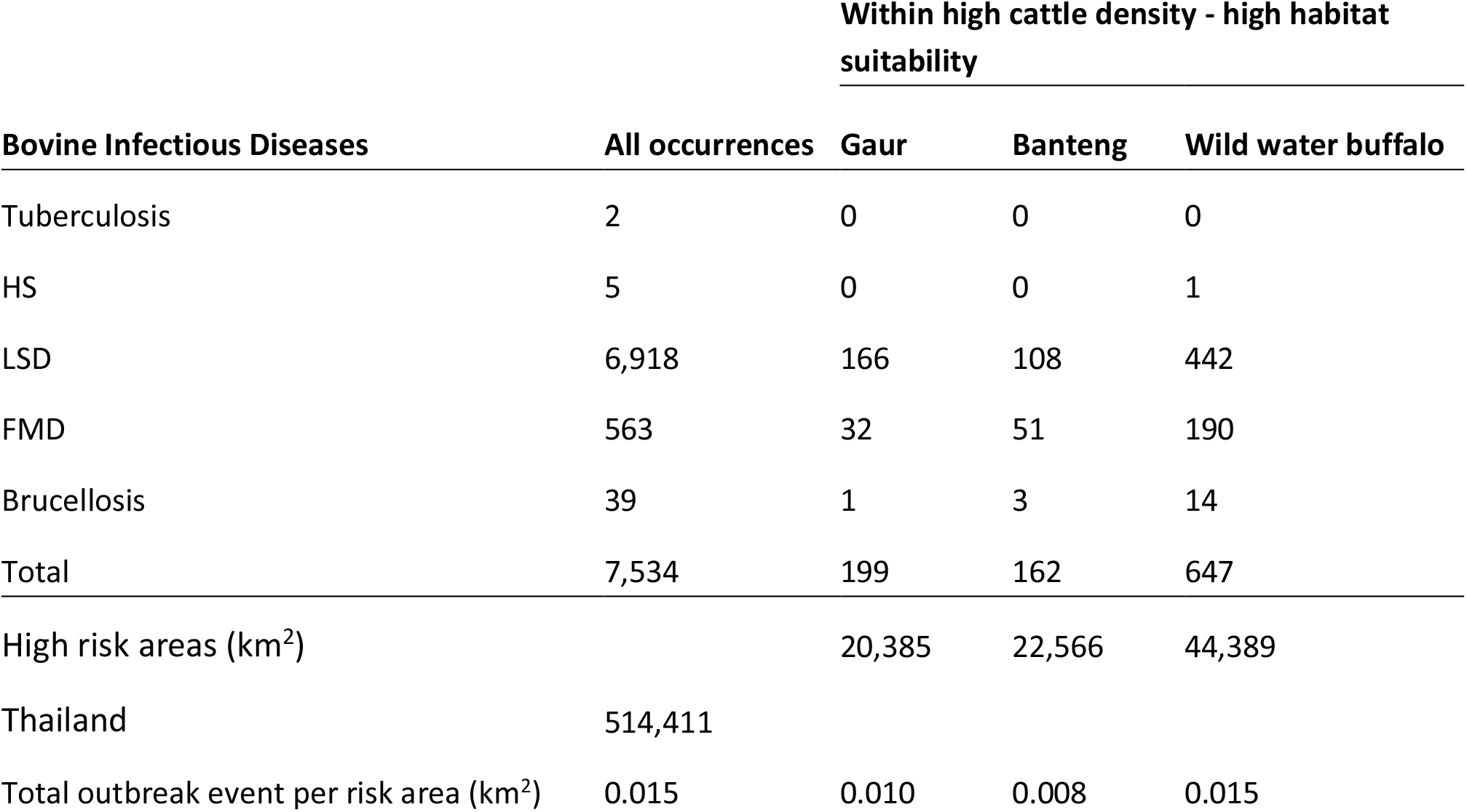
The reported outbreak events of bovine infectious disease occurrences in the high risk areas.

| Bovine Infectious Diseases | All occurrences | Within high cattle density - high habitat suitability |  |  |
| --- | --- | --- | --- | --- |
|  |  | Gaur | Banteng | Wild water buffalo |
| Tuberculosis | 2 | 0 | 0 | 0 |
| HS | 5 | 0 | 0 | 1 |
| LSD | 6,918 | 166 | 108 | 442 |
| FMD | 563 | 32 | 51 | 190 |
| Brucellosis | 39 | 1 | 3 | 14 |
| Total | 7,534 | 199 | 162 | 647 |
| High risk areas (km<sup>2</sup>) |  | 20,385 | 22,566 | 44,389 |
| Thailand | 514,411 |  |  |  |
| Total outbreak event per risk area (km<sup>2</sup>) | 0.015 | 0.010 | 0.008 | 0.015 |

Moreover, according to the land use types, the most extensive interface areas were found in close unknown forests (meaning they did not match any of the other forest definitions), followed by cropland for three species. Closed evergreen forests also contain risk areas for gaur and banteng. The open deciduous forest had no interface areas detected (Figure 3).

**Figure 3.**
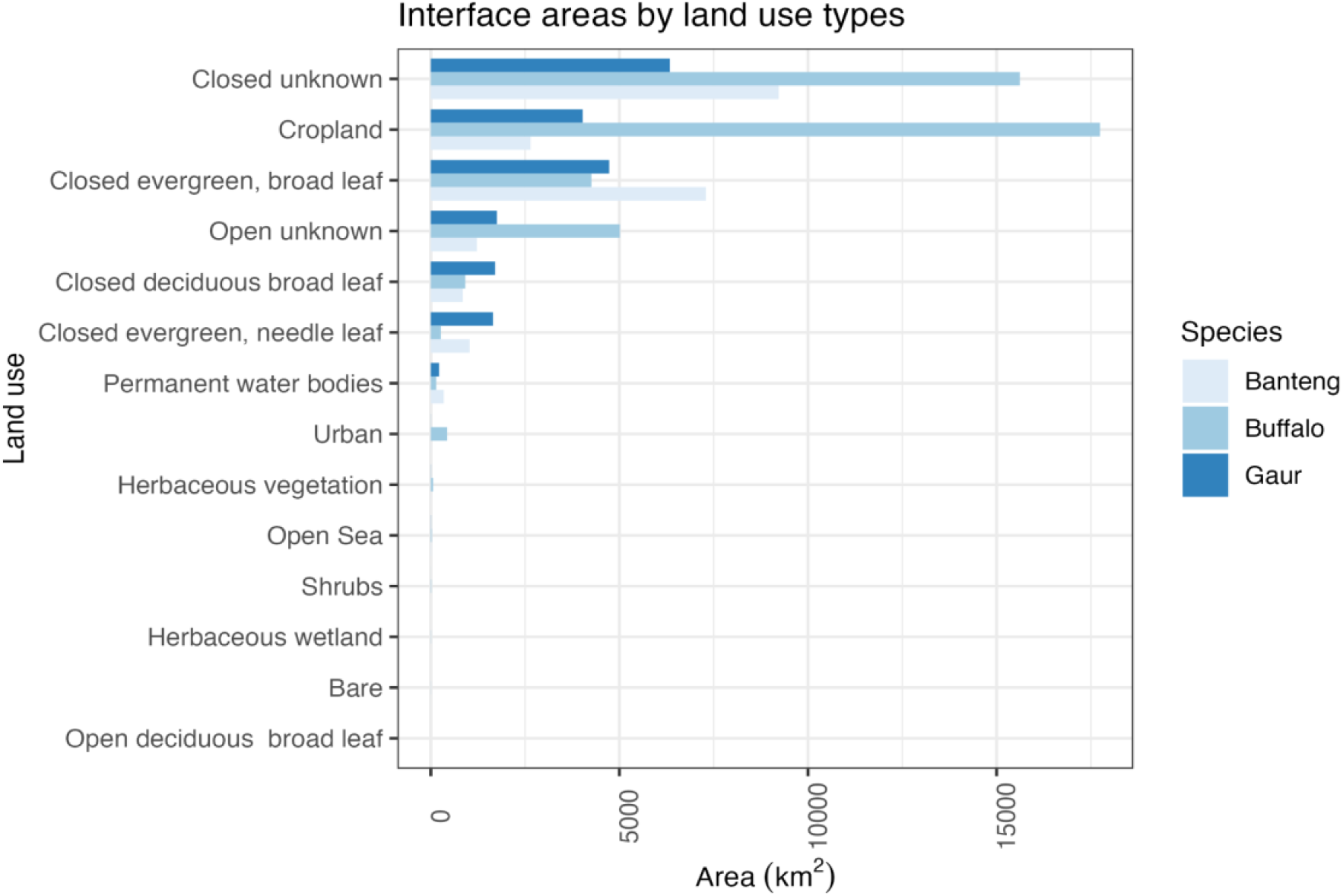
Interface areas of three wild bovid species are categorised based on land use types, from the land cover layers dataset ((41), https://zenodo.org/records/3243509). The term closed unknown forest denotes a type of forest that does not match any of the other definitions.

## Discussion

We examined potential risk areas of disease transmission between wild bovids and livestock and provided the preliminary focus area that should be considered for disease surveillance and mitigation in Thailand. Total risk areas in Thailand are between 4% (gaur and banteng) to 9% (wild water buffalo) of the country, with the most high risk areas being from the central west (Ratchaburi) to the south (Nakhon Si Thammarat). The highest risk proportion inside PAs was at Namtok Sam Lan National Park (NP) in the central, and the largest risk areas were Kuiburi NP and Kaengkrachan NP in the western forest, related to the highest cattle density in Thailand. Gaur and banteng have similar risk areas mostly around the edge of forests, while wild water buffalo have risk widely across the country because models predicted extensive amounts of suitable areas in the central down to the southern part that coincide with the high cattle density areas.

Although the wild water buffalo showed the largest areas of predicted suitable habitat among all three bovid species, it is also the most endangered. This is due to its current distribution being confined solely to the Huai Kha Khaeng Wildlife Sanctuary, with the population remaining stable and not exceeding 69 individuals for decades (42, 43). Thus, many areas identified as high risk are unlikely to be current high risk areas but are important for future planning should wild buffalo ranges expand to these or they be relocated. This species is highly susceptible to endemic infectious diseases that could rapidly lead to serious decline or even local extinction. For instance, diseases like HS can cause a high fatality rate of up to 80% in domestic buffalo (44). The highly contagious and fatal nature of diseases like rinderpest and FMD may be contributing factors to population disappearances in Nepal and India (45). An outbreak could also lead to local extinction in a single fragmented population, as the recovery process is prolonged and potentially results in a lack of gene flow (46), especially with the independent mother origin of Thai wild water buffalo (47).

In contrast to wild water buffalo, gaur and banteng have more opportunities for contact with domestic cattle and humans, while wild water buffalo may encounter livestock and humans encroaching into the protected areas and suitable habitat. Gaur and banteng can share habitats, making the interface areas similar to each other. However, gaur uses a wider range of habitat types (e.g., evergreen, deciduous dipterocarp, mixed deciduous forests) than banteng, which is restricted to dry and open forests (e.g., dry dipterocarp, mixed deciduous forest) (48, 49). These two species show evidence of contact with livestock and humans due to their ability to adapt and tolerate human activities, resulting in conflicts in overlapping areas (11, 12, 50). However, their habitat suitability decreases when the distance is closer to human settlement and the presence of domestic cattle grazing (51).

Our study identified the potential high-risk areas in the northeastern and southern parts, which have the highest cattle density but low or even an absence of the actual species distribution in some areas. This caveat is observed in the ecological niche modelling of wild water buffalo, where high habitat suitability represents potential distribution and may not necessarily correspond to the actual species distribution. Nevertheless, this caveat could be mitigated by collecting and regularly updating occurrences of these bovid species, as well as data on livestock density and distribution, or by restricting the analyses to areas with sufficient data.

Urbanisation and expansion of agricultural areas increase the opportunity for contact between domestic livestock and wildlife. Contact rate, the probability of transmission and the location shifts of animals at each time step, are the major factors that need to be considered for the spatial disease transmission model (52). The direct contact between wildlife and livestock is unlikely but indirect contact in the same space at different times via shared resources (e.g. water, grassland, supplement) with domestic animals potentially causes a chance of wildlife exposure to pathogens and disease transmission (13, 53). However, to succeed in cross-species infection, several factors should converge to drive pathogens through the natural barriers before having a pathogenic infection into a new host (54). We found that the highest interfaced areas were identified in cropland and unclassified forests, which potentially can be shared by free-ranging livestock and wild bovids. Thailand also experienced significant land cover changes (55) primarily driven by the increase of crop plantations and urbanisation with concurrent population growth, which leads to changing wildlife and livestock interactions and risk of disease transmission as per the previous studies (56).

Host density is one of the main risk factors in wildlife and livestock disease transmission (57, 58). We used host density and distribution as the main risk factors to identify and prioritise the potential risk areas of wildlife and livestock disease transmission, as has been used in the other studies (53, 59, 60). The advantage is knowing the target place for implementing the disease surveillance system, but the disadvantages can arise from the complexity of disease transmission dynamics, which depend on factors such as host species movement (60), contact pattern (61), high adaptability of wildlife behaviour, transmission modes (e.g. density or frequency-dependent) (58, 62) and population size (63).

Understanding the underlying factors that contribute to disease outbreaks in a specific potential risk area is essential for planning effective disease mitigation and control strategies. Multidisciplinary approaches incorporating key elements like pathogens, hosts, and environmental factors (supplementary materials Table S1-S2, describe the generic risk factors for disease transmission and mitigation methods between wildlife and livestock) also help policymakers develop disease control and mitigation measures. Pathogen spillover events are complicated, with a convergence of risk factors, which are difficult to approach. Integrating a complex system of human, animal and environmental will benefit prevention efforts or at least mitigate the impact of the next spill-over event (64).

Mitigation strategies will likely vary according to local socioeconomic conditions, but among the preventive actions are using vaccination of livestock or even wild species, targeted reduction of infected individual animals, herds or farms (ideally with compensation), along with reducing livestock herd sizes and densities, transport of livestock among farms, and contacts between farmed animals and wild species (65, 66). Contact reduction might be through measures such as altering land use at the local level, or with “natural” (e.g. plant-based) or artificial (e.g. metal) fencing or barriers and zoning of forests, livestock and human settlements to minimise the contact (17, 67), which may lead to reduce pathogen spillover (68). Longer term strategies might include societal transitions to lower meat-based, more plant-based proteins to reduce demand for meat and dairy products. Conserving intact forests with effective surveillance can mitigate the risk of disease transmission at the interface, especially in edge or transition areas. In contrast, fragmented forests increase the likelihood of wildlife being exposed to livestock and humans, leading to an elevated risk of disease transmission (20, 69, 70).

Livestock vaccination is crucial for reducing outbreak incidences of endemic diseases, requiring approximately 80% coverage to effectively prevent disease transmission for many pathogens based on pathogen specific, particularly in high-risk areas and populations, as part of routine practice (71, 72). However, capturing and delivering parenteral-route vaccinations to free-ranging wild bovids pose significant challenges, especially in tropical forests where animals might be hidden. Consequently, various aspects must be carefully considered in the vaccination plan, including the target population, coverage, safety, and efficiency, to effectively stimulate herd immunity (73). Non-invasive vaccination methods, like tuberculosis oral vaccination, have been tested in domestic cattle and some wildlife and are planned for use in wild cattle (74). Research and development for other endemic diseases like FMD, HS, and brucellosis is still ongoing (75-77). Culling livestock infected with zoonotic diseases (e.g., bTB, brucellosis) is commonly implemented in Thailand (78, 79). However, infected animals often undergo illegal translocation, potentially spreading the disease to other locations. To manage this issue, the government should rigorously regulate animal movement, regulating the guidelines for isolation of infectious animals during outbreaks, and providing appropriate compensation for culling cases. The effectiveness of these mitigation actions is influenced by the presence and use of effective infection and disease surveillance, as discussed below.

Livestock and, likely, wildlife disease surveillance in Thailand is based on the DLD, DNP (Thailand) and WOAH guidelines (80), which cover significant transboundary disease outbreaks in the country. Even though the passive surveillance system is useful for recording the obvious clinical signs and emerging infectious diseases (like FMD and LSD), there is a gap in collecting non-clinical to subclinical signs of disease due to these being challenging to detect. Moreover, passive surveillance leads to underreporting by the farmers for some zoonoses like bovine tuberculosis and brucellosis, for which animals must be condemned due to the slow and partial (not less than 75% of the market price, but often not 100%) compensation from the government. Another drawback is the clinical signs reported from passive surveillance may not refer to the place where the animal got infected if those are moved from the original area. Therefore, active surveillance such as risk-based (81), disease surveys (Arjkumpa et al., 2020) screening or detecting seroprevalence are necessary in hotspots or endemic areas to effectively allocate resources for disease mitigation and control strategies. Furthermore, even when reported, further work must be undertaken to understand the disease risk in depth. Reported data might refer to one event being a single case or multiple cases, and infectious diseases are, by default, dynamic in their nature. Without significant further work, passive surveillance data may offer a limited and biased understanding of the true disease risk in a location at a particular time. The impact of the detection of some infections on trade must be addressed, as this might be a barrier to effective surveillance and reporting (82, 83). The involvement of field authorities is another crucial aspect of data collection, indispensable for obtaining real-time information. Modern technologies or data sharing can help identify risk areas and plan preparedness implementations (62, 84, 85). Therefore, one should consider investing more in field data collection and incorporating field practitioners or epidemiologists into the team before formulating policies (86).

Using a One Health approach, a framework for disease surveillance has been developed, incorporating essential considerations of spillover events into the processes. To sustainably manage data collection and surveillance systems, collaboration among government organisations and stakeholders is a key step in the process, involving considerations of political, ethical, administrative, regulatory, and legal (PEARL) aspects through all approaches (87). An effective surveillance system, characterised by rapid detection and accurate results, not only monitors emerging diseases but also reduces the risk of disease transmission and minimizes the impact on lives, economies, and biodiversity during disease outbreaks (88). Also, the use of non-invasive data collection for willdife disease surveillance and surveys, such as faces, urine, saliva and environmental samples (e.g. soil, water) should be considered to avoid direct contact and reduce disturbing wildlife during capturing and data collection (89). Further studies may consider including other risk factors such as multi-species host distribution, the distance of risk factors and contact pattern (90, 91), as well as improving the model by using updated disease surveillance data and wild bovid species occurrences, especially for areas where the uncertainty of model predictions is high.

## Conclusion

Our study predicted the potential risk areas by using the interface areas between wildlife and domestic cattle, where livestock disease is frequently reported. We overlaid suitable habitats of three large wild bovids in Thailand with cattle density to create potential risk maps. High-risk areas were identified in locations with both high cattle density and high habitat suitability, particularly at the edges of forest-protected areas. Notably, small, fragmented forest areas with high cattle density presented the highest proportion of the high-risk areas. Among various land-use types, cropped land and some closed forests exhibited the largest interface areas. Our findings highlight the importance of wildlife habitat and intact forest conservation to mitigate contact and reduce vulnerability to extinction, reduce shared areas and address the potential risk areas for disease transmission between wild bovids and livestock. This methodology not only supports disease surveillance but also facilitates the implementation of effective mitigation and control measures.

## Acknowledgements

WH was supported by Manaaki New Zealand Scholarship. D.T.S.H., RSJ & RLM were supported by Bryce Carmine and Anne Carmine (née Percival), through the Massey University Foundation (grant no. RM22688), and D.T.S.H. was supported by the Percival Carmine Chair in Epidemiology and Public Health and Royal Society Te Apūrangi (grant no. RDF-MAU1701). Thank you to Dr Weerapong Thanapngtham from DLD, Thailand, for sharing the disease surveillance data.

## Supplementary Materials

Table S1: Generic risk factors for disease transmission in wildlife and livestock interface areas, including studies and perspectives in Thailand and internationally. Table S2: Mitigation methods for infectious disease transmission.

